# Characterization of Dependencies Between Growth and Division in Budding Yeast

**DOI:** 10.1101/082735

**Authors:** Michael B. Mayhew, Edwin S. Iversen, Alexander J. Hartemink

## Abstract

Cell growth and division are processes vital to the proliferation and development of life. Coordination between these two processes has been recognized for decades in a variety of organisms. In the budding yeast *Saccharomyces cerevisiae*, this coordination or ‘size control’ appears as an inverse correlation between cell size and the rate of cell-cycle progression, routinely observed in G_1_ prior to cell division commitment. Beyond this point, cells are presumed to complete S/G_2_/M at similar rates and in a size-independent manner. As such, studies of dependence between growth and division have focused on G_1_. Moreover, coordination between growth and division has commonly been analyzed *within* the cycle of a single cell without accounting for correlations in growth and division characteristics *between* cycles of related cells. In a comprehensive analysis of three published time-lapse microscopy datasets, we analyze both intra-and inter-cycle dependencies between growth and division, revisiting assumptions about the coordination between these two processes. Interestingly, we find evidence (1) that S/G_2_/M durations are systematically longer in daughters than in mothers, (2) of dependencies between S/G_2_/M and size at budding that echo the classical G_1_ dependencies, and, (3) in contrast with recent bacterial studies, of negative dependencies between size at birth and size accumulated during the cell cycle. In addition, we develop a novel hierarchical model to uncover inter-cycle dependencies, and we find evidence for such dependencies in cells growing in sugar-poor environments. Our analysis highlights the need for experimentalists and modelers to account for new sources of cell-to-cell variation in growth and division, and our model provides a formal statistical framework for the continued study of dependencies between biological processes.

## I. Introduction

Cell division and cell growth are processes fundamental to all life, and their dysregulation is common in diseases like cancer. In the budding yeast *Saccharomyces cerevisiae*, cell division is known to be coordinated with cell growth [1–6] (reviewed in [7]). This dependence between growth and division is most noticeable in daughter cells that, owing to the asymmetric manner of budding yeast division, are born smaller than their mothers (Figure 1). Consequently, daughters undergo longer G_1_ phases to reach a ‘critical size’ at START, the point of cell-cycle commitment [8, 9]. Generally, a correlation has been observed between the birth mass of a cell and its time spent in G_1_, with smaller cells at birth taking longer to complete G_1_. This ‘size control’ is important for the maintenance of a consistent size distribution in the cell population from generation to generation (size homeostasis).

**FIG. 1:**
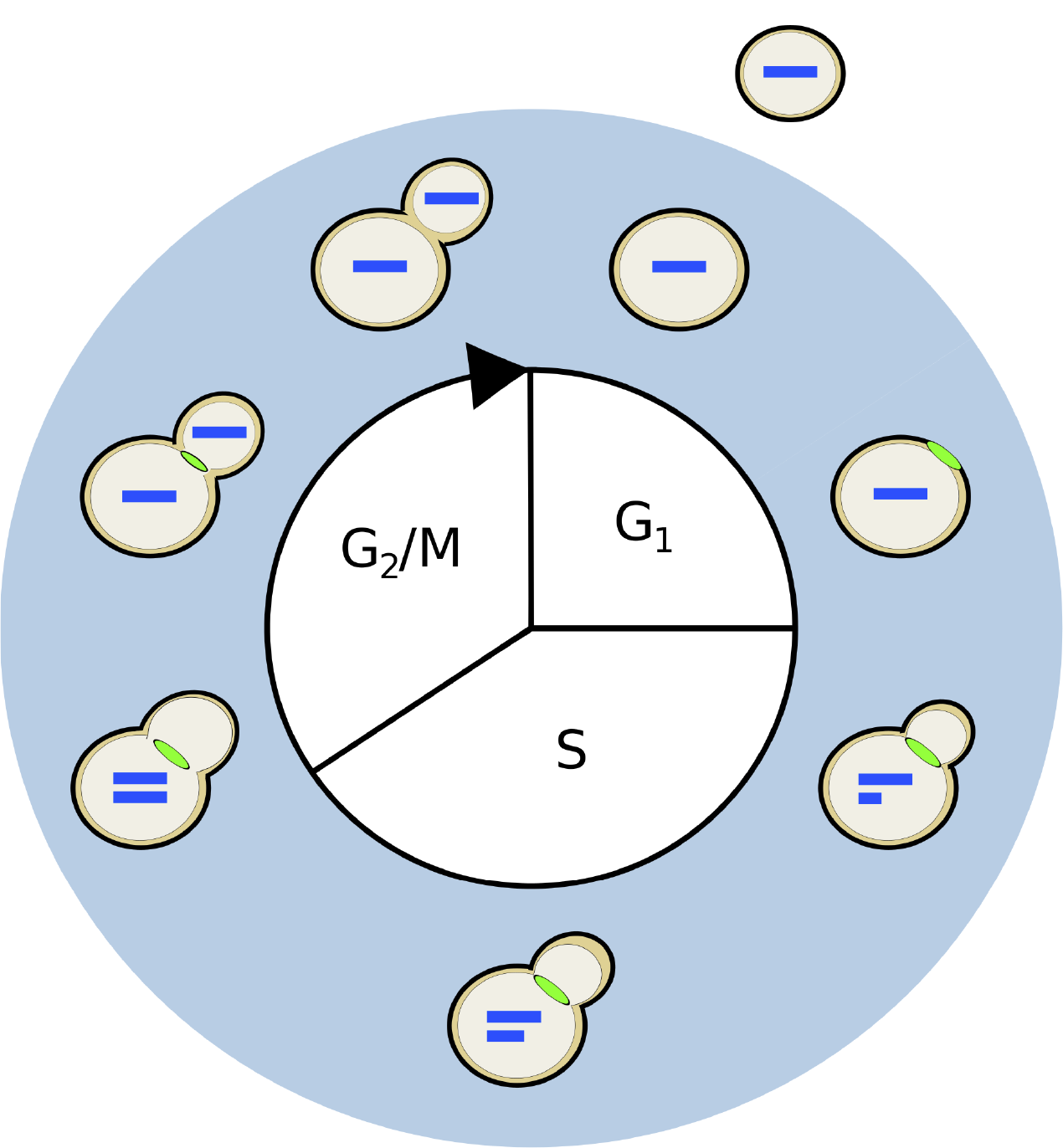
A diagram of the haploid budding yeast cell cycle. The process of cell division begins at the top of the diagram and proceeds clockwise. Cell division consists of four phases: G_1_, S, G_2_, and M; the latter two have been merged since classical studies demonstrate that they largely overlap in yeast. Along the outer ring of the diagram are depicted progressive stages of division, as reflected by the different markers of cell-cycle progression. Each bar inside the cell represents a single copy of DNA. The feature at the neck joining the mother and daughter cell represents the myosin ring. The myosin ring appears late in G_1_, marking the location where the bud will emerge, and disappears with cytokinesis, indicating the separation of the mother and daughter cytoplasms. After cell wall separation, the mother and daughter cells are free to undergo more rounds of division. In budding yeast, division is asymmetric and daughters (shown outside) are born smaller than their mothers.

Studies of coordination between growth and division in budding yeast have focused primarily on G_1_. Indeed, after having reached a sufficient and roughly similar size, both mother and daughter cells are presumed to proceed through the S, G_2_, and M phases at similar rates [9, 10]. These tenets of the traditional model of budding yeast size control set certain expectations for S/G_2_/M duration: (1) G_1_ duration is size-dependent, while S/G_2_/M duration is size-independent, and (2) S/G_2_/M duration is roughly constant from cell to cell. Assuming a consistent single-cell growth rate across cells, these tenets are sufficient to maintain a consistent size distribution in the population. Recent studies in bacteria have revealed an alternative size control model by which cells add a relatively constant amount of volume over the cell cycle, regardless of their birth size [11–13]. Recent time-lapse microscopy datasets tracking characteristics of individual cells provide evidence with which to test these models and further characterize coordination between growth and division in budding yeast.

These time-lapse datasets also allow investigation of correlations *between* measurements made at different cell cycles, an important gap in our understanding of coordination between growth and division. In multicellular systems, coordination of division among cells has important implications for higher-scale phenomena like development, differentiation, and tissue organization [14–17]. In unicellular organisms like the budding yeast, *Saccharomyces cerevisiae*, inter-cycle dependencies between growth and division are plausible [18] and might affect more classically studied intra-cycle dependencies or characteristics. For example, a mother cell with some advantage in cell division or growth might transmit that advantage to her progeny, resulting in a fast-dividing or fast-growing daughter cell. However, it is unclear the extent to which cell-cycle progression is correlated across cell cycles in budding yeast, if at all.

Statistical modeling provides a powerful and principled foundation for characterizing these correlations in lineages of proliferating cells. Indeed, correlation and biological lineage analysis have been intertwined since the development of the correlation coefficient by Galton and Pearson [14, 18–22]. Statistical models of correlation in cellular characteristics have been successfully applied to bacterial and mammalian cell lineage data [23, 24]. In addition, modern Bayesian inference techniques for regression and model averaging provide a framework for evaluating the plausibility of a variety of different models of correlation between growth and division [25].

Here, we provide in-depth statistical analysis to address four main biological questions: (1) Is S/G_2_/M duration approximately constant across cells or does it vary among mothers and daughters?; (2) Is S/G_2_/M duration independent of cell size?; (3) Is size at birth independent of size accumulated over the cell cycle?; and (4) Is there evidence for inter-cycle dependencies in cell-cycle progression that accompany known intra-cycle dependencies? We analyze three microscopy datasets comprising different genetic and nutrient environment conditions [2]. We conduct a Bayesian regression analysis to further investigate the dependence between cell growth and division within and between cycles and comprehensively evaluate the plausibility of different models of correlation. We introduce a novel hierarchical statistical model [26, 27] of budding yeast cell division at the single-cell level to formally characterize inter-cycle correlations in cell-cycle progression. Our analysis offers fresh biological and methodological insights on the extent and nature of coordination between cell division and cell growth, as well as a novel framework for formally characterizing dependencies within and between cells in these and other biological processes.

## II. Single-Cell Analysis of Budding Yeast size Control Models

### A. Single-cell measurements of *Saccharomyces cerevisiae* growth and division

Single-cell data of haploid budding yeast were acquired from a previously published study [2]. The study followed cell-cycle progression and growth in 26 wild-type lineages (782 cells) grown in glucose, 19 6×CLN3 lineages (376 cells) grown in glucose, and 21 wild-type lineages (518 cells) grown in glycerol/ethanol. Only those cells (or a subset thereof where specified) with fully observed cell-cycle durations were retained for subsequent processing and analysis, resulting in 213 wild-type cells in glucose, 99 6×CLN3 cells, and 157 wild-type cells in glycerol/ethanol.

Cell-cycle progression was measured using the times of occurrence of two landmark cell-cycle events for each yeast cell on the plate: the appearance and disappearance of the myosin ring, visualized by tagging Myo1p with green fluorescent protein (GFP) (Figure 1). The myosin ring is a contractile structure that appears late in G_1_, just prior to the appearance of the bud [28] (Figure 1). Cell growth was monitored with the red fluorescent protein, DsRed, which was placed under the control of the promoter of ACT1, the constitutively expressed actin gene. In this way, total red fluorescence in a cell served as a proxy for total protein content or cell mass. Red fluorescence in a cell was quantified at each time point and suitably normalized across all cells in a microcolony (Supplement).

### B. Size-dependent differences in S/G_2_/M duration between mothers and daughters

Budding yeast cells born at a smaller than average size tend to undergo longer G_1_ phases to reach a sufficient size for cell-cycle entry, manifesting as a dependence between cell size and division within a given cell cycle. Under the assumption that mothers and daughters complete G_1_ at roughly the same size, the cells could maintain a consistent cell size distribution from generation to generation provided the amount of time they spent in S/G_2_/M was similar on average. As such, we tested whether combined S/G_2_/M duration is similar across mother and daughter cells [9].

To assess differences in the observed mother and daughter S/G_2_/M duration, we performed two-sided nonparametric Wilcoxon rank sum tests in the three datasets (Figure 2 and Table I). For this analysis, we separated mother and daughter S/G_2_/M durations. We used a subset of the mother cells since some S/G_2_/M durations were associated with the same mother in consecutive cell cycles (Supplement). We found significantly shorter S/G_2_/M’s for mothers compared with daughters for both 6×CLN3 cells and wild-type cells growing in glucose. We also found suggestive (but not significant) differences in S/G_2_/M for wild-type cells growing in glycerol/ethanol.

**FIG. 2:**
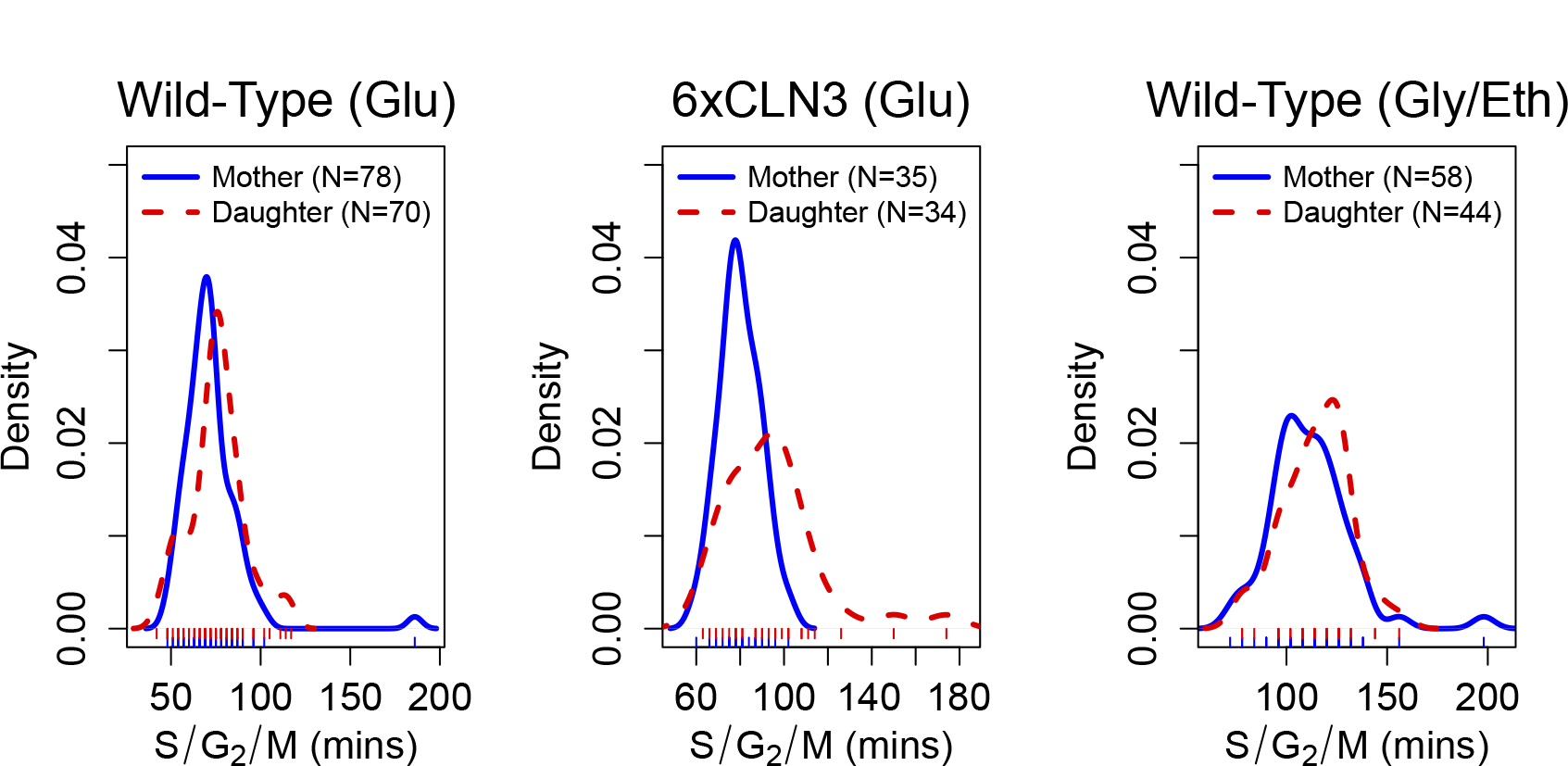
Density plots of S/G_2_/M durations for mother and daughter cells in the three different experimental conditions. Rug plots appear below each density plot.

**TABLE I:** Differences in S/G_2_/M duration between mother and daughter cells. Cell counts are the same as in Figure 2. Estimates of differences in S/G_2_/M duration and 95% confidence intervals (in parentheses) are shown. *Before rounding of interval estimates for display purposes, interval for wild-type cells in glycerol/ethanol included 0 while interval for glucose did not (hence different p-values).

| Dataset | Estimate (mins) | P-value |
| --- | --- | --- |
| Wild-Type (Glu) | -6 (-9,0) | 0.021 |
| 6×CLN3 | -12 (-18,-3) | 0.001 |
| Wild-Type (Gly/Eth) | -6 (-12,0)* | 0.166 |

One potential explanation for these differences in S/G_2_/M duration is that daughter cells smaller than their mothers at the onset of S/G_2_/M might require more time to complete the budded period. We tested this hypothesis by pairing mother cells from the previous analysis with their first daughters. Here, mother-daughter pairs only consisted of daughter cells whose immediate mother was not also a daughter cell herself. We then fit a linear regression of the daughter-mother difference in log S/G_2_/M duration on the differences in estimated mass at budding (Figure 3). In this regression model, the intercept represents the conditional expected difference between daughter and mother log S/G_2_/M durations independent of size differences and the coefficient is the slope in terms of differences in fitted masses at budding. In two of the three experimental settings, the difference between daughter and mother mass at budding was a significant negative predictor of differences in log S/G_2_/M duration (Table II). Thus, daughters smaller than their mothers at the point of cell-cycle entry tended to spend more time in S/G_2_/M. Collectively, these results provide evidence that S/G_2_/M duration is systematically different between mother and daughter cells, and, surprisingly, we identify a growth-related component to this difference.

**FIG. 3:**
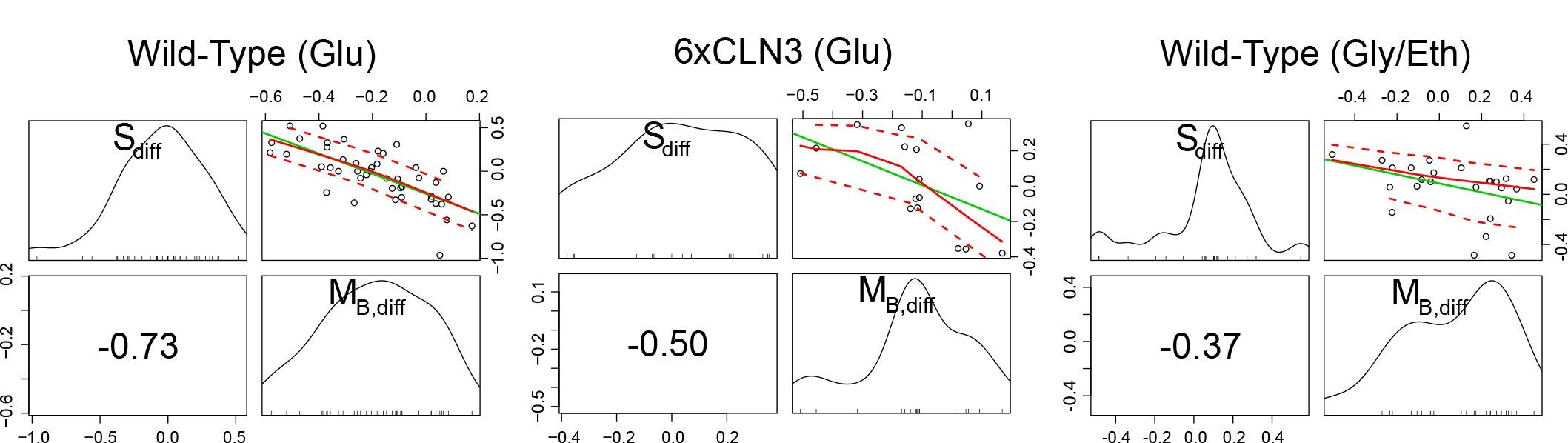
Scatter and marginal plots of mother-daughter differences in log S/G_2_/M duration and mass at budding. Marginal density plots of each variable are shown on the diagonal. In the lower off-diagonal panels appear the Spearman rank correlation coefficients. Scatter plots along with best linear fit lines (green), loess smoothed fit lines (solid red) and loess spread lines (dashed red; root-mean-squared positive and negative residuals) appear in the upper diagonal panels. The loess span was 1.0. Daughter-mother pair counts are those listed in Table II.

**TABLE II:**
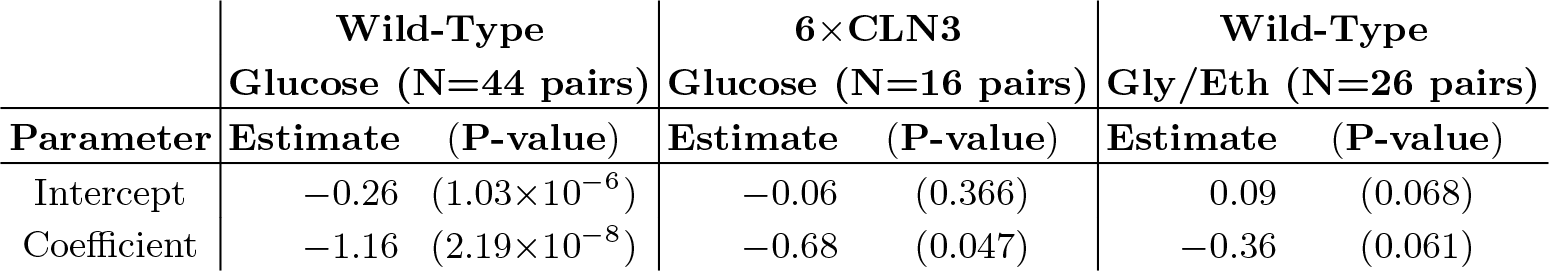
Regression of daughter-mother differences in size at budding on differences in log S/G_2_/M duration. P-values (in parentheses) are for the test of zero-valued estimates. Fitted masses at budding were computed from linear regressions on the logarithm of each cell’s growth traces (Section II, Supplement). P-values less than 0.001 are shown in scientific notation.

|  | Wild-Type |  | 6×CLN3 |  | Wild-Type |  |
| --- | --- | --- | --- | --- | --- | --- |
|  | Glucose (N=44 pairs) |  | Glucose (N=16 pairs) |  | Gly/Eth (N=26 pairs) |  |
| Parameter | Estimate | (P-value) | Estimate | (P-value) | Estimate | (P-value) |
| Intercept | -0.26 | (1.03×10<sup>-6</sup>) | -0.06 | (0.366) | 0.09 | (0.068) |
| Coefficient | -1.16 | (2.19×10<sup>-8</sup>) | -0.68 | (0.047) | -0.36 | (0.061) |

### C. Significant associations observed between size at birth and size accumulated

Recent studies in bacteria and budding yeast have suggested a compelling alternative model for size control called the ‘adder’ model [6, 11–13] which posits that the size added by cells during the cell cycle is roughly constant from cell to cell and independent of the cell’s size at birth. To evaluate evidence for this hypothesis in our data, we analyzed the association between the observed birth mass (M_0_) and mass accumulated between birth and division (M*_add_* = M*_div_* – M_0_). Interestingly, we find strong negative correlations between mass at birth and mass accumulated over the cell’s life in every cell type and every condition (see Figure 4). Moreover, the correlations we observe are all significant with two-sided hypothesis tests of the Spearman rank coefficients (Table III).

**FIG. 4:**
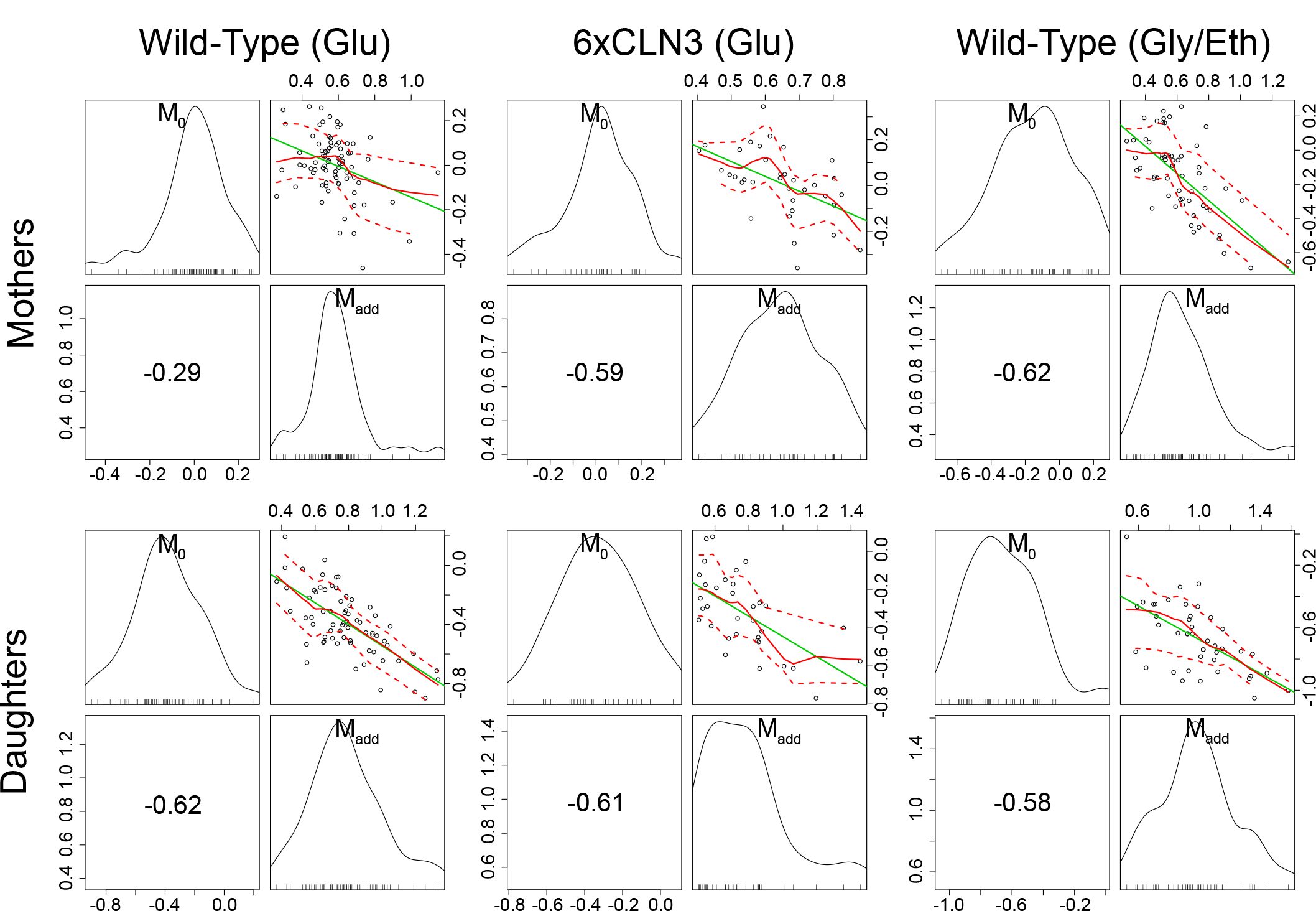
Scatter plots and univariate density plots of mass at birth and mass accumulated from birth to division for mother and daughter cells in all three experimental conditions. The layout and content of each of the six panels is the same as in Figure 3. Cell counts are in Table III.

**TABLE III:** Test of no correlation between size at birth and mass accumulated over the cell cycle. The third column shows p-values for the correlation between mass at birth (M_0_) and mass added over the entire cell cycle (M_*add,full*_). The fourth column shows p-values for the correlation between birth mass and mass added during S/G_2_/M (M_*add,sg2m*_). P-values less than 0.001 are shown in scientific notation.

| Dataset | Cell Type | $M_{add,full}$<br>P-value | $M_{add,sg2m}$<br>P-value |
| --- | --- | --- | --- |
| Wild-Type (Glu) | Mothers (N=78) | 0.011 | 0.188 |
| | Daughters (N=70) | $2.10 \times 10^{-8}$ | 0.183 |
| 6×CLN3 | Mothers (N=35) | $2.22 \times 10^{-4}$ | 0.003 |
| | Daughters (N=34) | $1.69 \times 10^{-4}$ | 0.001 |
| Wild-Type (Gly/Eth) | Mothers (N=58) | $3.82 \times 10^{-7}$ | 0.001 |
| | Daughters (N=44) | $4.49 \times 10^{-5}$ | 0.172 |

One possible explanation for the association we observe is that it is driven primarily by a negative correlation between mass at birth and size accumulated during G_1_ (more classical size control dependence) and that mass at birth and size accumulated during S/G_2_/M are uncorrelated. However, we observe significant negative associations between mass at birth and size accumulated during S/G_2_/M, particularly in 6×CLN3 cells (Table III). These correlations might indicate a compensatory mechanism during S/G_2_/M to overcome disabled G_1_ size control and ensure robust cell size at division. In aggregate, we find no evidence for ‘adder’ model effects in our time-lapse datasets.

### D. Post-G_1_ dependence between cell-cycle progression and cell growth

As aforementioned, budding yeast daughter cells tend to spend more time in G_1_ than their mothers to reach a sufficient size for cell-cycle entry. This reflects an association between G_1_ duration and cell size at birth. It has been hypothesized that G_1_ is the primary period during which cell-cycle progression depends on cell size and that S/G_2_/M progression is largely independent of size, subject instead to a timing mechanism [10]. Moreover, analyses of coordination between growth and division have focused primarily on dependencies *within* rather than *across* cell cycles. However, given that budding yeast cells divide asymmetrically, leading to partitioning of organelles and other cellular contents between mothers and daughters, it is plausible that cell-cycle progression might depend on characteristics of the cell’s mother as well as on the size of the cell itself.

Classically, one would analyze the correlation between a cell-cycle interval (e.g. G_1_) and the cell’s size at the beginning of that interval. Here, by conditioning on more predictor variables, we can estimate the relative effects of a cell’s size and the growth and division characteristics of its mother on the cell’s current cell-cycle durations. So, we first computed growth characteristics of a cell and its immediate antecedent cell. Using the single-cell growth traces of each cell *j* and its immediate predecessor cell (*Pa*(*j*); mother cycle that immediately preceded cycle of cell *j*) in lineage *i*, we estimated growth-related variables (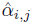, 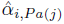, 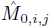, 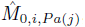, 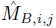, and 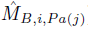) by assuming exponential single-cell growth kinetics and fitting a separate linear model to the logarithm of each cell’s growth trace ([2]; Supplement). The fitted intercept of this linear model gave the estimated birth mass 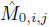 of cell *j* from each lineage *i* while the slope gave the estimated mass accumulation rate (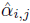). We also retained the fitted mass at budding of each cell (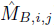). We then fit linear regression models of log S/G_2_/M durations on these cell-level estimates as well as on the log S/G_2_/M durations of the cell’s predecessor (*S*_*i, Pa*(*j*)_).

Here, a model represents a particular pattern of dependence between growth and division and is determined by the set of predictor variables included in the regression. As we have 7 different predictor variables (not counting the included intercept term, *μ_S_*), there are 2^7^ = 128 possible regression models. To infer the most plausible model of dependence between size and cell-cycle progression while explicitly accounting for uncertainty in the model specification, we conducted Bayesian model averaging [25]. Since we didn’t have strong prior information about the dependencies between these variables, we assumed that each regression model was equally plausible *a priori*. We then computed posterior probabilities of each model for mother cells (Figure 5) and daughter cells (Figure 6). For the analysis of mothers, we retained every other cell cycle of the mother cell starting with its most recent cycle in the lineage, accounting for potential correlations between different cycles of the same mother. This procedure resulted in 53 wild-type pairs in glucose, 25 6×CLN3 pairs, and 46 wild-type pairs in glycerol/ethanol. For the analysis of daughters, we used the mother-daughter pairs of a previous analysis (cell counts provided in Table II).

**FIG. 5:**
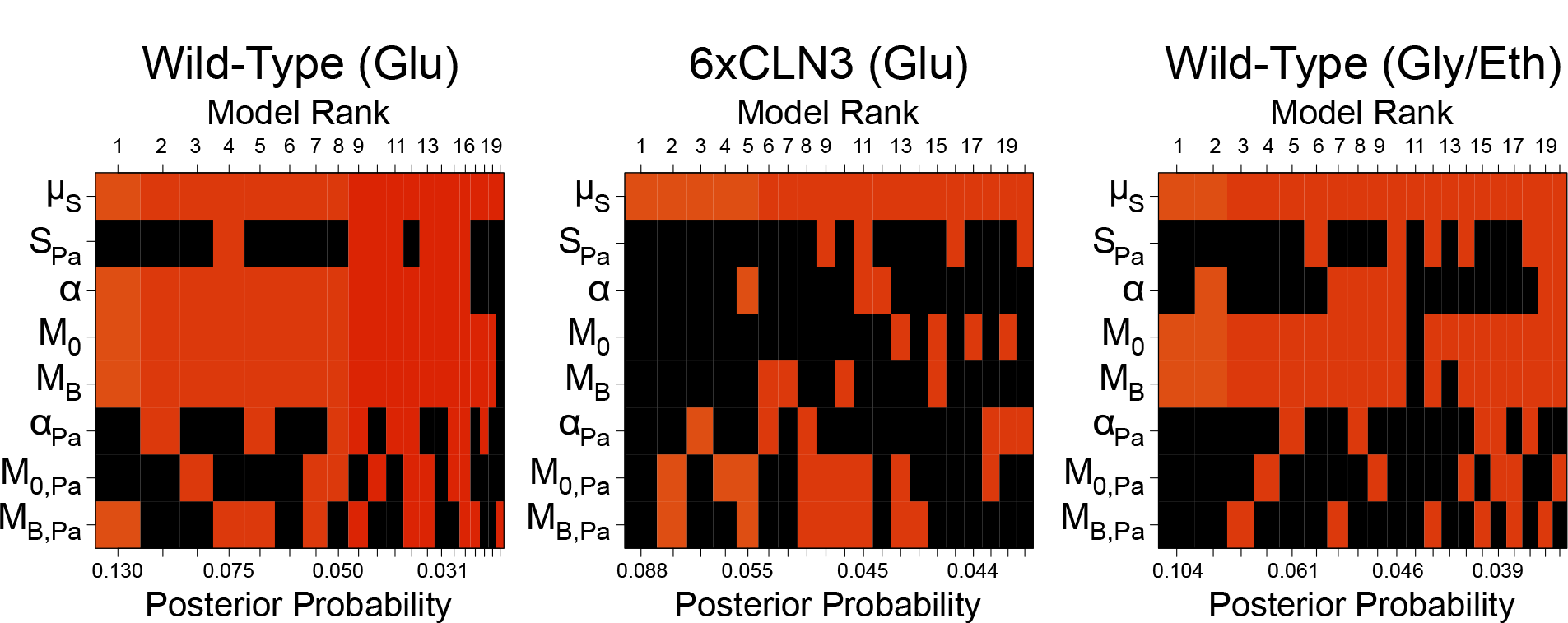
Bayesian adaptive sampling results for mother cells. Shown are the top 20 models (columns ranked by posterior probability) where colored squares indicate that the corresponding predictor variable is included in the model and black squares indicate exclusion of the variable.

**FIG. 6:**
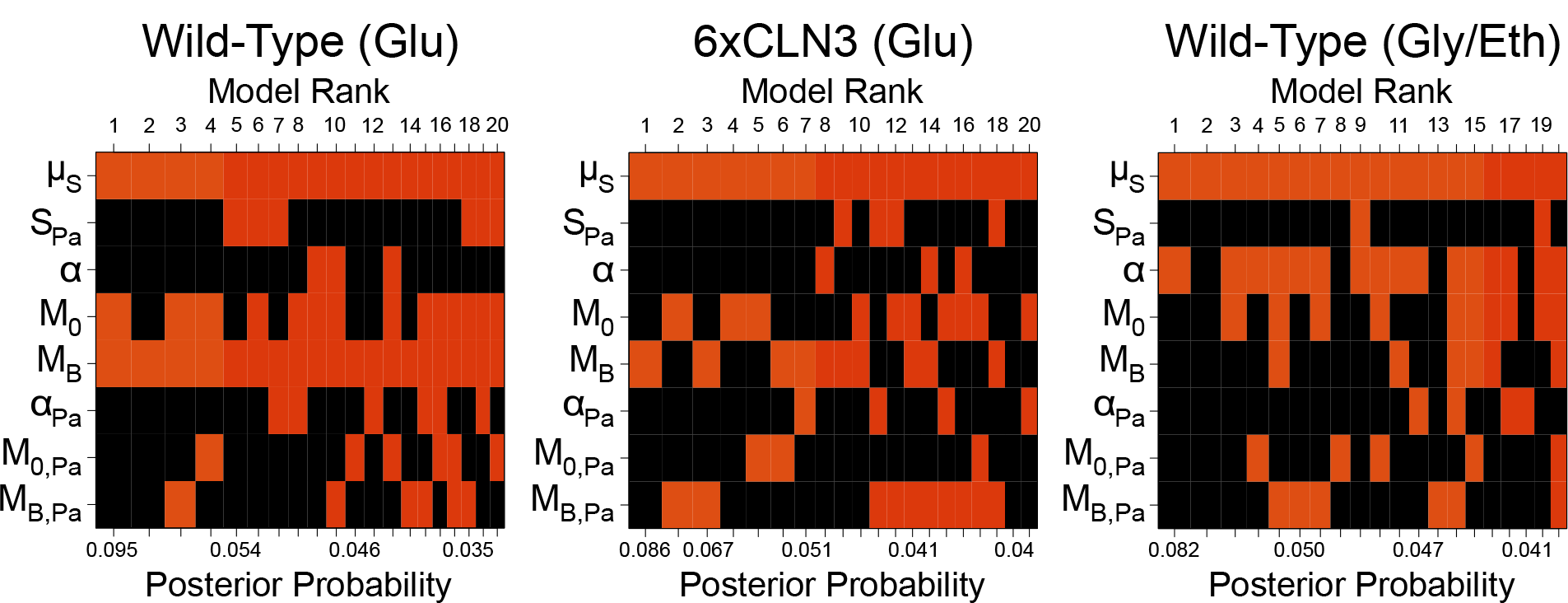
Bayesian adaptive sampling results for daughter cells. Panel layout and content is the same as in Figure 5.

When we consider mother-to-mother dependencies, we find a strong association for wild-type mothers between log S/G_2_/M duration and growth characteristics, particularly mass at budding (Figure 5, left and right panels). Mass at budding (*M_B_*) was included in nearly every enumerated model with non-zero posterior probability (Figure 5) indicating that mass at budding was an informative predictor of mother log S/G_2_/M duration (˜82% posterior probability of inclusion; log_10_(Bayes factor) = 0.657 or ‘substantial evidence’ for inclusion [29]). Mass at birth (*M*_0_; ˜74% posterior probability) and mass accumulation rate (*α*; ∼71% posterior probability) also tended to be included as predictors. Likewise, for a mother growing in glycerol/ethanol, we detect an association between her current log S/G_2_/M duration and mass at birth (∼64% posterior probability) and budding (∼61% posterior probability; Figure 5, right panel). In addition, the posterior means (averaged across all models) of the included regression coefficients for mass at budding for wild-type mothers in glucose (–1.129) and glycerol/ethanol (–0.262) were consistent with classical G_1_ size control (larger mass at budding corresponds to less time spent in S/G_2_/M). This pattern of dependence has not previously been observed in mother cells, potentially due to the fact that we are conditioning on multiple growth and division characteristics for the cell’s current and previous cycles. We did not see such patterns of dependence for 6×CLN3 mother cells. Importantly, we also did not find strong evidence for dependence between a wild-type or 6×CLN3 mother’s log S/G_2_/M duration and her characteristics in her previous cycle (Figures 5). So, conditioned on summaries of the mother’s current cycle, her log S/G_2_/M duration can be considered independent of her previous growth and cell-cycle progression.

Extending this analysis to mother-to-daughter associations, we again discovered patterns of dependence between a cell’s log S/G_2_/M duration and mass at budding (*M_B_*): for wild-type daughter cells growing in glucose (Figure 6, left panel). Mass at budding was included as an explanatory variable for the wild-type daughter’s log S/G_2_/M duration (in glucose) in nearly all models with non-zero probability (∼93% posterior probability of inclusion; log_10_(Bayes factor) = 1.118 or ‘strong evidence’ for inclusion according to [29]; Figure 6, left panel). As in the previous analysis, the model-averaged posterior mean of the included regression coefficient was –0.616, an estimate consistent with classical G1 size control. We also found mild associations between log S/G_2_/M duration and other growth characteristics in all three conditions (Figure 6). In particular, we note an association between log S/G_2_/M duration and mass accumulation rate in glycerol/ethanol (∼58% posterior probability of inclusion). The model-averaged posterior mean of the included regression coefficient for mass accumulation rate was –18.391, indicating that daughters with larger mass accumulation rates spent less time in S/G_2_/M. Collectively, our findings for both mother and daughter cells run counter to the notion that S/G_2_/M duration is independent of size.

## III. Hierarchical Modeling of Correlation in Budding Yeast cell Division at the Single-Cell Level

The regression framework used in the previous section highlighted associations between growth and division within and across cycles. However, we limited our analysis to rigidly defined mother-mother and mother-daughter pairs and did not take advantage of the inherent hierarchical organization of the data (i.e. cells make up lineages, multiple lineages are observed for each experimental condition). Moreover, simply computing sample-based estimates of inter-cell correlations would preclude separation of cell-to-cell variation in cell-cycle progression from variation due to measurement error. Hierarchical models provide a formal framework to represent such structure and naturally pool information across replicate lineages as well as allow for estimation of cell-specific and noise-related sources of variation. An important property of these models is the potential to reduce parameter estimation error relative to sample-based approaches by shrinking cell-specific parameter estimates towards sample (population) estimates [26]. For these reasons, and to more effectively characterize dependencies between cells, we opted to analyze single-cell cell-cycle durations with a novel hierarchical model.

### A. Observing a cellular branching process

To motivate our model, we consider how the single-cell data are acquired using time-lapse microscopy. A single yeast cell (the founder cell; cell 1 in Figure 7) growing on the agarose slab is identified at the onset of the time-lapse experiment. Each founder cell generates a lineage consisting of two fully observed sub-lineages (Figure 7). The mother sub-lineage consists of the founder’s first cell cycle after division from its daughter (‘mother origin’) and her progeny. The daughter sub-lineage consists of the founder cell’s first daughter (‘daughter origin’) and her progeny. Non-origin cells of these sub-lineages are hereafter referred to as ‘mother’ and ‘daughter’ cells.

**FIG. 7:**
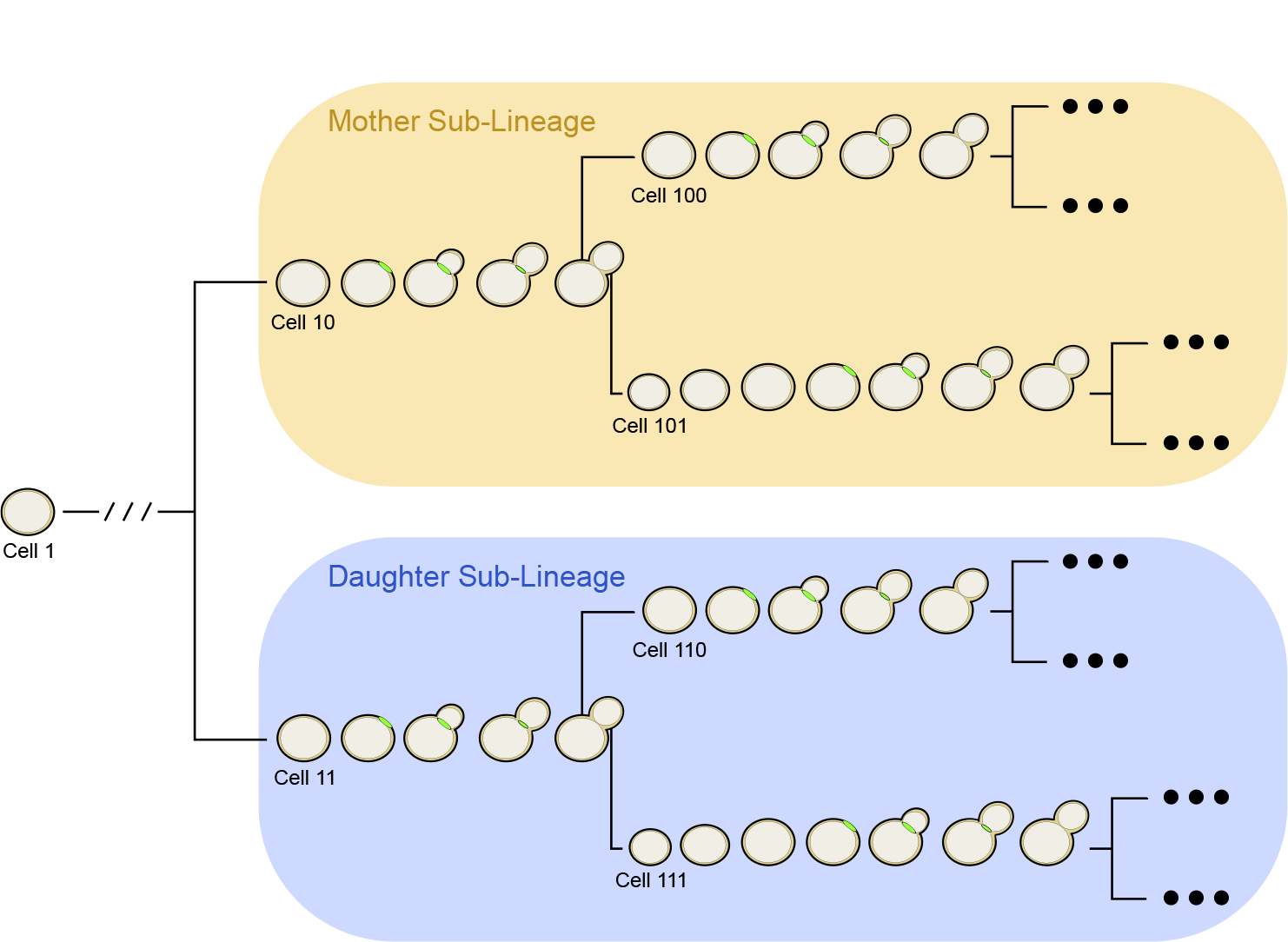
Illustration of single-cell lineages and classification of cell types. Shown is a typical single-cell lineage tree from the dataset of Di Talia *et al.*, 2007. Arranged along each branch of the lineage tree are images of representative cells undergoing cell-cycle events. Binary cell labels ending in 0 indicate a mother cycle while labels ending in 1 indicate a daughter cycle.

For these sub-lineage cells, we observe the time since the start of the time-lapse experiment of the appearance and disappearance of the cell’s myosin ring. We refer to these times as budding and division times, respectively. In our notation, the budding time for cell *j* from lineage *i* is *B_i,j_* and the division or cycle time is *C_i,j_* (Figure 8). We transformed these times to cell-specific budding (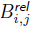) and division (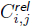) durations (Figure 8; Supplement). To refer to durations specific to each cell, we adopted the binary indexing scheme of Di Talia et al. [2].

**FIG. 8:**
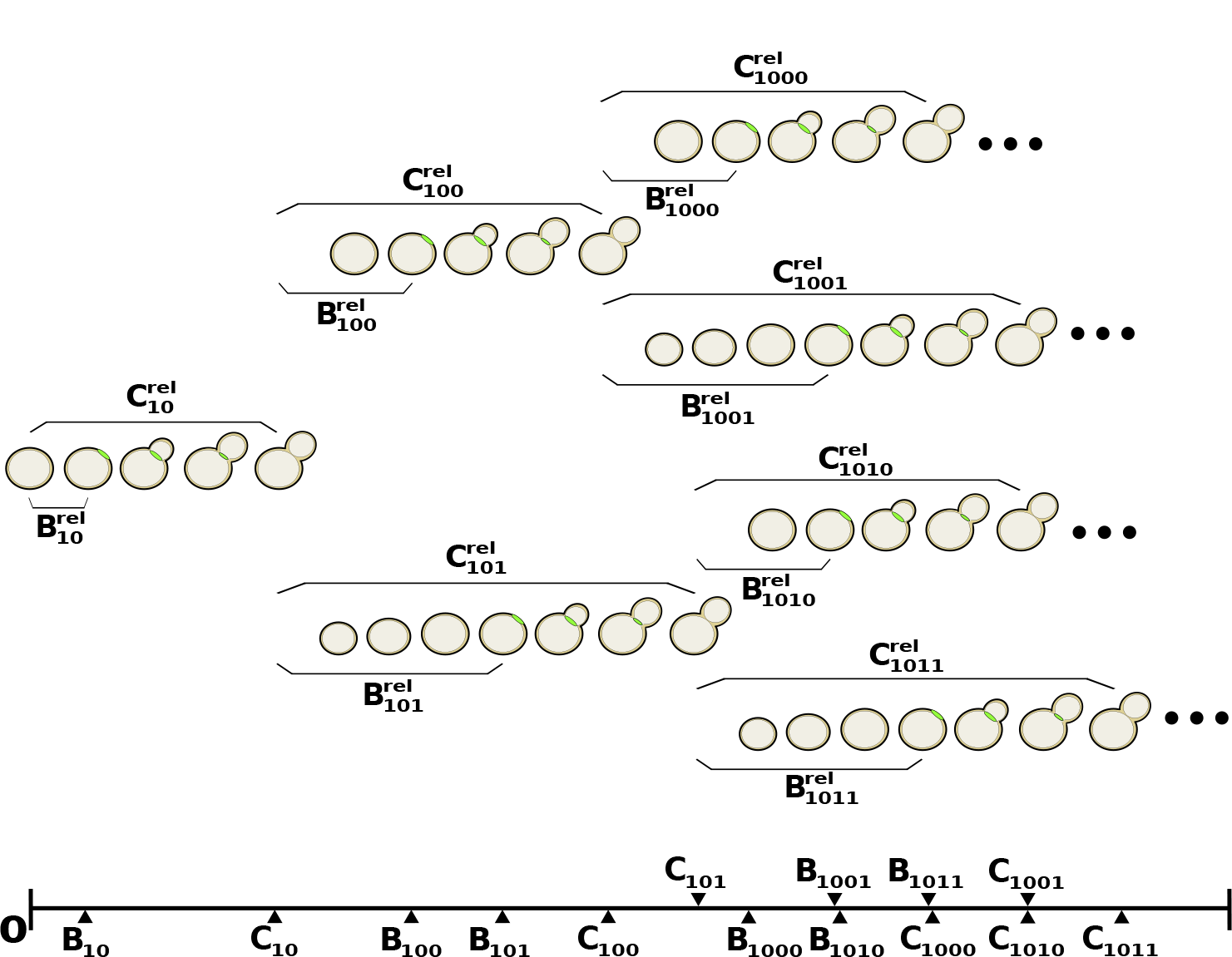
Sample diagram of budding and division observations arising from the time-lapse microscopy experiments. Here, an example mother origin cell (cell 10) of the sub-lineage proceeds through the cell cycle, undergoing budding and division. Dividing from her daughter (cell 101), the mother origin becomes mother cell 100 and undergoes another round of division. The ellipses following the lineage leaves indicate that other lineages could be larger.

We view the budding and division durations from each lineage (as in Figure 8) as noisy observations from an underlying branching process (illustrated in Figure 9). The branches of the tree in Figure 9 represent the expected observed division durations for each cell. The cell’s expected budding duration is some fraction of this branch length. To identify inter-cycle correlation in cell-cycle progression, we assume that the branch lengths in the process may depend on one another. Further details of model construction follow.

**FIG. 9:**
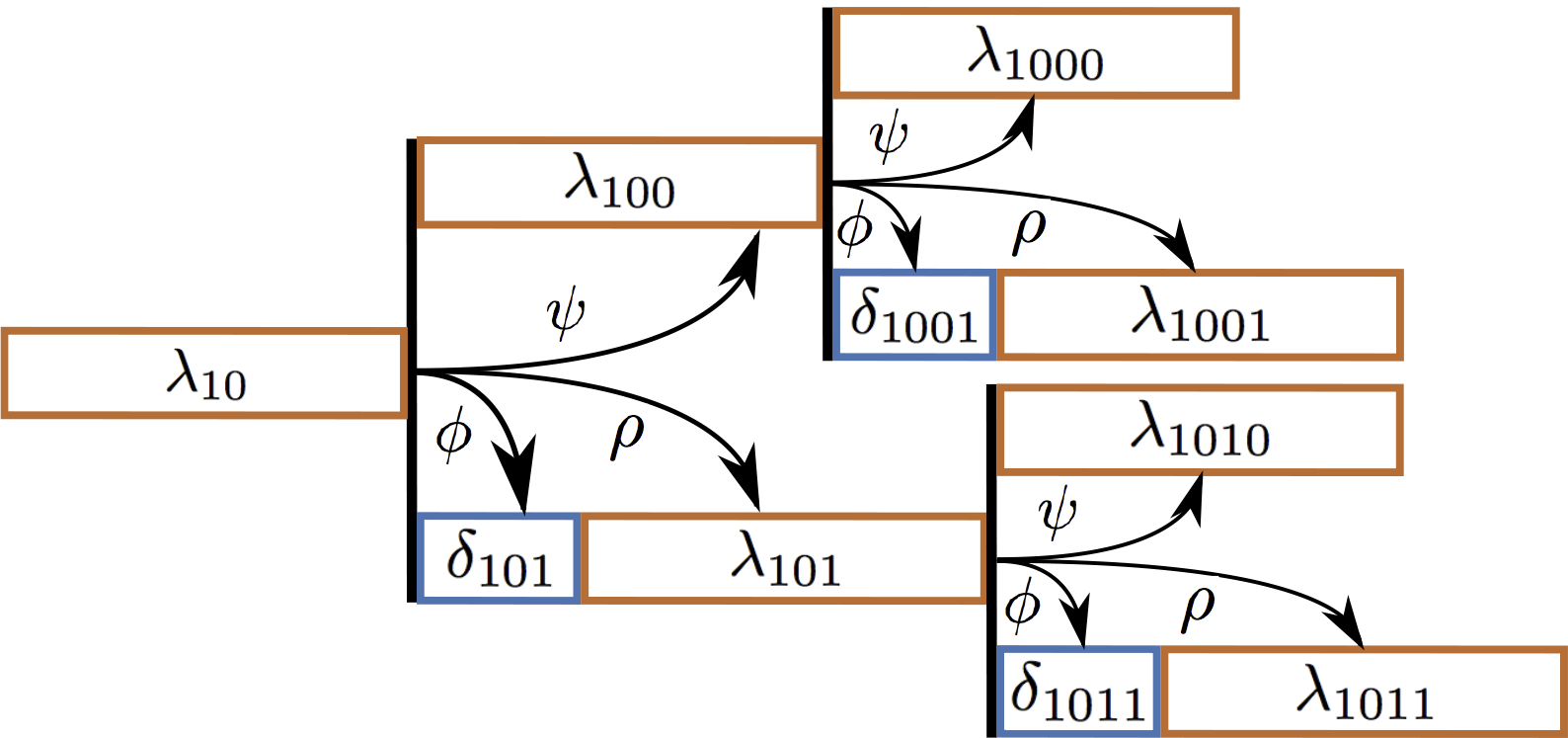
A diagram of the asymmetric branching process specifying expected cell-cycle durations for each cell. The diagram is drawn to indicate the branch lengths that give rise to the budding and division durations in Figure 8. λ_10_ is the expected cell-cycle duration of mother origin cell 10. The expected cell-cycle duration of her subsequent cycle is λ_100_, which depends on her first cell cycle through the correlation parameter *ψ*. For the daughter branch, two parameters specify the cell’s expected division duration: *δ*_101_ and λ_101_. These branch lengths depend on the mother’s cell-cycle duration through the correlation parameters ρ and *ϕ*, respectively.

### B. Likelihood and error model for budding and division observations

The first (or lowest) level of our hierarchical model captures noise or error in observations while higher levels of the hierarchical model will capture cell-to-cell variability. Here, we first assumed that the elapsed times from which the durations for cell *j* in lineage *i* were derived were independent and normally distributed with means *μ_B__i,j_* and *μ_C__i,j_* and variance *τ*^2^. While the observed cell-cycle durations are positively valued, we use normal errors at the lowest level of the hierarchical model rather than alternative error models (e.g. log-normal) as we don’t expect multiplicative errors from the manual recording of budding and division events. Rather, as will be discussed in a later section, we constrain the branch length parameters to be positively valued. After transformation of elapsed times into durations, the likelihood of all budding and division durations for lineage *i* is:

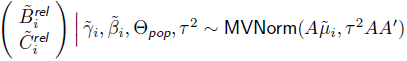

Here, *A* is a linear transformation matrix and 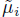 is a vector of the expected budding and division durations for lineage *i*. 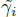 is a vector containing two types of cell-specific parameters that make up the branching process of a lineage: the expected base cell-cycle durations for each cell in lineage *i* (*λ_i,j_*s) and extensions to expected daughter cell-cycle durations due to smaller size at birth (*δ_i,j_*s). Θ*_pop_* is the set of population-level parameters {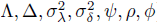} (see Table IV) that specify correlation and cell-to-cell variation in the *λ_i,j_*s and *δ_i,j_*s. The vector 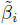 contains another set of cell-specific parameters (*β_i,j_*s): the fraction of a cell’s *λ* branch in the unbudded phase. We describe the model for the means (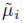) of the observations and the remaining parameters in the next section.

**TABLE IV:** Population parameters of hierarchical model

| Parameter | Description |
| --- | --- |
| $\Lambda$ | population average of mother cell-cycle duration (minutes) |
| $\Delta$ | population average of daughter cell G<sub>1</sub> extension duration (minutes) |
| $\psi$ | correlation between λs from two successive mother cycles |
| $\rho$ | correlation between a mother's $\lambda$ and her daughter's $\lambda$ |
| $\phi$ | correlation between a mother's $\lambda$ and her daughter's $\delta$ |
| $\sigma_\lambda^2$ | variance in cell-specific $\lambda$ branch lengths (minutes<sup>2</sup>) |
| $\sigma_\delta^2$ | variance in cell-specific $\delta$ branch lengths (minutes<sup>2</sup>) |
| $\mu_m$ | expected proportion of $\lambda$ spent unbudded in mothers |
| $\mu_d$ | expected proportion of $\lambda$ spent unbudded in daughters |
| $\eta_m$ | prior weight of information for mother unbudded period |
| $\eta_d$ | prior weight of information for daughter unbudded period |
| $\tau^2$ | variance in measurement error (minutes<sup>2</sup>) |

### C. Representing inter-cycle dependence and cell-to-cell variability with an asymmetric branching process

The second level of the hierarchical model consists of cell-specific parameters (e.g. branch lengths) that give the expected value of a cell’s budding and division durations for a particular lineage. This second level comprises a branching process in which the branch lengths are potentially correlated and vary from cell to cell (Figure 9). The branch lengths for a lineage *i* are represented by a vector 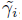, composed of two sets of parameters: *λ_i,j_*s and *δ_i,j_*s. The *δ*s account for longer daughter cell cycles due to smaller birth sizes. To measure correlation between these branch lengths, we introduced three parameters: *ψ*, *ρ*, and *ϕ* (Figure 9, Table IV)

As noted in previous work [30, 31] and since the cell-cycle durations are positively valued, we jointly model all branch lengths for a lineage *i* (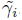) with a multivariate log-normal distribution. That is:

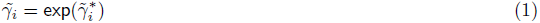

with

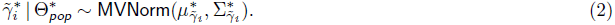

Here, 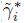 is the vector of cell-specific branch lengths on the natural logarithmic scale. These log-scale durations follow a multivariate normal distribution with a structured covariance matrix 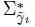. In this covariance matrix, we encode a simple model of inter-cycle dependence. With no strong expectations of the extent of inter-cycle correlation structure, we consider the simplest model for inter-cell correlation: that the expected log-scale cell-cycle duration of a newly arisen cell depends solely on the expected log-scale cell-cycle duration of its predecessors in the lineage only through its mother. As we do not observe the cell-cycle durations of each lineage’s founder cell (Figure 7), this correlation structure dictates that the branch lengths of the mother origin (*λ_i,j_*) and daughter origin are correlated with one another, and we model them accordingly (Supplement).

The mean vector 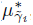 consists of parameters Λ^*^ and Δ^*^ (counterparts of Λ and Δ on the log scale). 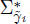 is parameterized by log-scale analogues of *ψ*, *ρ*, *ϕ*, *σ_λ_*, and *σ_δ_*. To infer parameters on the original scale (Table IV), we transform the log-scale analogues (Supplement).

As with the *λ* and *δ* parameters, each cell has a parameter indicating the proportion of its *λ_i,j_* it spends in the unbudded state: *β_i,j_*. The vector 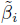 comprises these cell-specific parameters for lineage *i*.

Now, we can compute the expected value of a cell’s observed budding and division durations. For example, if a cell *j* is a mother cell then:

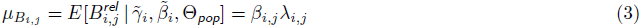

and

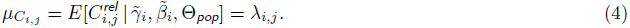

On the other hand, if cell *j* is a daughter cell, then:

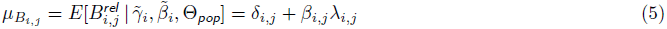

and

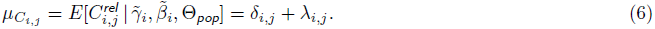

The third level of our hierarchical model (represented by population-level parameters Θ*_pop_*; Table IV) encapsulates patterns of cell-cycle progression and inter-cycle dependence shared across replicate lineages in a given experimental condition. Owing to the hierarchical structure of the data, we assume for each experimental condition that the branch lengths (*λ_i,j_*s and *δ_i,j_*s) from all lineages are drawn from the same distribution: the multivariate log-normal distribution parameterized by Λ, Δ and other parameters of Θ*_pop_*. More formally, the branch lengths are exchangeable within a given experimental condition [26]. We make a similar assumption for the cell-specific *β_i,j_*s: the unbudded proportions are exchangeable within a given cell type (mother or daughter) and experimental setting.

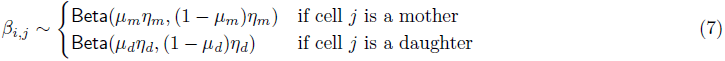

### D. Hierarchical model fitting uncovers variation in cell-cycle progression across experimental conditions and between mothers and daughters

Fits of the hierarchical model to the three different datasets suggest distinct patterns of cell-cycle progression. As shown in Table V, population average mother cell-cycle duration (Λ) was approximately 88 minutes for wild-type cells in glucose. In contrast, mother cells divided nearly twice as slowly in glycerol/ethanol (∼146 minutes). Since Cln3 is a rate-limiting factor for cell-cycle entry [2, 32], daughters with 6 copies of CLN3 show quite short G_1_ extensions (*δ*) compared to wild-type daughters grown in glucose (Table V). Likewise, the estimated spread in 6×CLN3 daughter G_1_ extensions (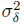) was much smaller compared to the corresponding estimates for wild-type cells, reflecting greater availability of Cln3 protein [2]. In contrast, wild-type daughters in glycerol/ethanol took nearly 95 minutes more to complete G_1_ (on average) than their mothers [33].

**TABLE V:** Posterior inferences (modes and 95% highest posterior density intervals) for model parameters. Λ, ∆, *σ_δ_*, *σ_λ_*, and *τ* are in minutes. Cell-specific unbudded duration parameters, *μ_m_* and *μ_d_*, range from 0 to 1.

|  | Wild-Type<br>Glucose (N=213) | 6×CLN3<br>Glucose (N=99) | Wild-Type<br>Gly/Eth (N=157) |
| --- | --- | --- | --- |
| Parameter | Estimate | Estimate | Estimate |
| $\Lambda$ | 87.60 (85.08,90.33) | 95.80 (92.40,100.79) | 145.24 (139.19,152.34) |
| $\Delta$ | 25.55 (20.21,31.30) | 18.57 (12.20,23.03) | 94.02 (81.68,107.78) |
| $\mu_m$ | 0.18 (0.16,0.19) | 0.15 (0.13,0.17) | 0.25 (0.23,0.27) |
| $\mu_d$ | 0.13 (0.09,0.17) | 0.05 (0.01,0.11) | 0.22 (0.17,0.27) |
| $\eta_m$ | 23.19 (18.45,28.77) | 25.48 (19.02,33.19) | 22.84 (17.89,28.49) |
| $\eta_d$ | 16.53 (11.51,23.33) | 20.39 (13.48,28.22) | 23.37 (16.56,31.56) |
| $\psi$ | -0.07 (-0.31,0.20) | -0.03 (-0.38,0.50) | 0.46 (0.22,0.65) |
| $\rho$ | -0.18 (-0.41,0.08) | -0.29 (-0.61,0.12) | 0.21 (-0.13,0.51) |
| $\phi$ | -0.20 (-0.40,0.05) | -0.64 (-0.84,0.01) | -0.02 (-0.26,0.24) |
| $\sigma_\delta$ | 20.00 (14.63,30.32) | 6.56 (2.83,10.11) | 42.65 (33.16,56.56) |
| $\sigma_\lambda$ | 15.30 (13.66,17.20) | 15.72 (13.49,18.82) | 26.07 (22.54,30.11) |
| $\tau$ | 1.31 (0.93,1.91) | 1.43 (0.99,2.30) | 1.45 (0.96,2.30) |

A benefit of our hierarchical approach is the ability to separate cell-to-cell variation in cell-cycle progression from measurement error. As mentioned previously, while inferences for *σ_λ_* do not change much between the two types of cells grown in glucose, *σ_δ_* is dramatically reduced in 6×CLN3 cells reflecting differences in cell-cycle progression one might expect. However, as the cells were grown in similar conditions, differences in cell-cycle progression should have little effect on an experimenter’s ability to record budding and division times, and so, measurement error should be similar. Importantly, inferences for *τ* for all three conditions are similar to one another (95% confidence intervals overlap).

As part of our deeper investigation of inter-cycle dependence, we generated inferences for the three correlation parameters in the model (*ρ*, *ψ*, and *ϕ*). As shown in Table V, no strong correlations exist in the two strains grown in glucose (wild-type and 6×CLN3) with all 95% posterior confidence intervals overlapping 0. However, we do see moderate mother-to-mother *λ* correlations (*ψ*) for wild-type cells grown in glycerol/ethanol. As we did not detect mother-to-daughter correlations in the same conditions, the inferences for *ψ* suggest that cells in glycerol/ethanol tend to retain the rate of cell-cycle progression with which they are born. This correlation could not be explained by drift in cell-cycle progression due to a cell’s replicative age or time spent by the cells on the plate (Supplement). However, considering our previous Bayesian regression analysis (Part I, Section D), this cell-cycle dependence is likely mediated by growth characteristics of the mother cell in her current cycle. To rule out the possibility that this result is an artifact of over-fitting the data, we carried out a leave-one-out cross-validation analysis to evaluate the capacity of different models of inter-cycle correlation to predict the observed cell-cycle durations of left-out cells. The results of this analysis were consistent with our parameter inferences in that models lacking *ψ* predicted the observed cell-cycle duration of wild-type cells in glycerol/ethanol more poorly (Supplement).

Overall, this statistical framework has generated valuable insights into potential inter-cycle sources of variation in biological processes.

## IV. Discussion & Conclusions

In this novel analysis we set out to address questions regarding dependencies within and between the fundamental processes of growth and division. We have found evidence from our analysis contradicting two tenets of budding yeast size control: a relatively constant S/G_2_/M duration shared by mother and daughter cells and a lack of dependence between S/G_2_/M duration and size. Our statistical analysis, including inferences from our hierarchical model (Supplement), also demonstrated that combined S/G_2_/M duration appeared longer on average in daughter cells compared with mother cells. Moreover, we detect a size-related component underlying these differences in S/G_2_/M duration. In support of our results, at least one classical study with single cells has noted that the budded duration was mildly longer (5–8 minutes on average) for daughters compared with mothers under a range of different growth conditions [34]. These observations are important because experimenters and modelers might otherwise assume approximately similar S/G_2_/M durations across cell types or simpler dependence structure between size, G_1_ and S/G_2_/M that might not be present in their experimental conditions, potentially affecting downstream conclusions about coordination between growth and division.

Our Bayesian regression analysis uncovered patterns of dependence between cell size characteristics and S/G_2_/M duration for wild-type cells in two different nutrient conditions. Post-G_1_ dependence between growth and division has been observed in budding yeast strains engineered for phase locking [35] and predicted by dynamic models of cell-cycle progression [31, 35]. In particular, we note phenotypic similarities between the post-G_1_ size dependence we observe in 6×CLN3 cells and that observed in a study of strains in which the G_1_ cyclin, CLN2, was under the control of the inducible MET3 promoter [35]. As in that study, we see both reduced G_1_ duration variability (Supplement; *σ_δ_* in Table V) and overall cell-cycle durations in 6×CLN3 cells similar to wild-type cells. Our analysis more comprehensively demonstrates these patterns in S/G_2_/M, identifying these dependencies in wild-type cells and in mother cells as well as daughter cells, on which size control studies have traditionally been focused. Our analysis also takes into account new sources of variation (e.g. inter-cell dependencies) to qualify these dependencies. This analysis coupled with our finding of no evidence for ‘adder’ model effects in our data add to evidence for compensation in cell-cycle time during S/G_2_/M and raises the question of whether size control exists outside of G_1_ in budding yeast.

Dual, complementary mechanisms of size control have been noted in the fission yeast, *Schizosaccharyomyces pombe*, with a strong size control imposed at the G_2_/M boundary and a weaker compensatory size control imposed at the G_1_/S boundary [10]. However, while our findings seem at first glance to extend the classical model of size control in budding yeast, we caution that this observed dependence does not necessarily imply a true size control mechanism. Rather, this association could be related to compensation in cell-cycle time due to premature cell-cycle entry or the activation of a cell-cycle checkpoint [36] due to perturbed cell-cycle progression.

Consequently, our analysis has generated experimentally testable hypotheses about the molecular basis of post-G_1_ size dependence and insights for future studies of size control. While the dependence we observe in 6×CLN3 cells, for example, is likely not due to activation of the morphogenesis checkpoint [37], other molecular targets related to DNA replication checkpoints (e.g. Rad53) or cryptic budding yeast size control (e.g. Bck2) should be tested to ascertain their relative effects on dependence between mass at budding and S/G_2_/M. Recent experimental work in budding yeast suggests an intriguing mechanistic model for size control in which dilution of Whi5 as the cell grows in volume dictates cell cycle entry at the G_1_/S transition [5]. This work could be extended to analyze changes in concentration of Whi5 and other proteins during S/G_2_/M to identify potential mechanistic bases for the dependencies we observe.

We do not detect evidence in our datasets for an ‘adder’ model of size control. Instead, we detect substantial negative dependencies between size at birth and size accumulated over the cell cycle. An important distinction between the current study and previous analyses is the measurement of cell size. In our datasets, cell size was measured via a fluorescent protein-based proxy for cell mass whereas recent work in bacteria and budding yeast has focused on cell volume [6, 11–13]. Elements of cell volume in budding yeast, particularly the vacuoles, are known to undergo dynamic, regulated changes over the course of the cell cycle [38]. On the other hand, it’s unclear the extent to which the fluorescent protein construct used in Di Talia’s datasets is the best proxy for cell size without direct measurements of cell mass. Therefore, further experimental and analytical studies are required to reconcile these results and determine whether the type of cell size measurement has any effect on mechanistic conclusions about size control.

In addition to these contributions, we have developed a novel hierarchical model of budding yeast cell-cycle progression. In uncovering correlations in cell-cycle progression between wild-type mother cycles in glycerol/ethanol, our model highlights a potential need for considering *between* as well as *within* cell dependencies. If capturing cell-to-cell dependence is an experimental goal, then time-lapse microscopy is preferable over techniques involving fixed and independent samples taken over time from an initially synchronized population of cells. From a statistical perspective, observing more lineages and more generations per lineage allows for better characterization of this dependence. Our analysis also suggests that both single-cell experimentalists and modelers should at least consider the possibility of such dependence to avoid potential confounding of other observed between-cell or within-cell dependencies. Dependence between cell growth and cell division has been thoroughly studied *within* a given cell cycle. However, the possibility that this correlation might be mediated by inter-cycle dependencies brought about by changes in environment or nutrient availability cannot be ignored. Correlations within a cycle could disappear or decrease in magnitude when conditioning on characteristics of the previous cell or generation. Conversely, conditioning on additional variables from previous cell cycles might not affect an observed correlation, providing greater context for experimental follow-up or model construction. In either case, our analysis demonstrates that both experimentalists and modelers can benefit from considering multi-generational data acquisition and analysis to verify the robustness of their correlation inferences.

Our model provides a flexible and extensible platform for analysis of intra-cycle as well as inter-cycle dependencies. The hierarchical specification of our model and our Bayesian approach to inference easily accommodates new lineage information. We also note that the model is not limited to cell division observations and can be adapted to the statistical analysis of dependence in any biological process (e.g. by dropping the budding yeast-specific *δ* and *ϕ* parameters and treating the *λ* branch parameters as the sole quantity of interest). In addition, while we made use of the single-cell growth data in our regression analysis, we are finalizing development of extensions to the hierarchical model to formally fit both the growth and division measurements in a joint analysis, making for a powerful tool to estimate correlations between multiple biological processes while accounting for dependencies between cells in a lineage.

This work represents an important step towards understanding the dependencies in cell-cycle progression and cell growth within and across cells in a dividing population. The statistical model-based approaches described here—coupled with ongoing time-lapse microscopy studies—will shed new light on cell-cycle and cell growth regulation and reveal mechanistic insights about the coordination between these two fundamental biological processes.

## V. Acknowledgments

We would like to thank Stefano Di Talia and Frederick Cross; Sung Sik Lee and Matthias Heinemann for generously providing their data; and Merlise Clyde, Stefano Di Talia, Steve Haase, Daniel Lew, Nick Buchler, Bruce Futcher, Kurt Schmoller, Ivan Surovtsev, members of the Haase and Hartemink labs, and the anonymous reviewers for comments about the analysis and the manuscript. This work was funded in part by grants from NIH (P50 30 GM081883-01) and DARPA (HR0011-09-1-0040 to A.J.H.) and was performed in part under the auspices of the U.S. Department of Energy by Lawrence Livermore National Laboratory under contract DE-AC52-07NA27344 (LLNL-JRNL-702334-DRAFT).

